# Wnt3 expression as a readout of tissue stretching during *Hydra* regeneration

**DOI:** 10.1101/2020.12.22.423911

**Authors:** Jaroslav Ferenc, Panagiotis Papasaikas, Jacqueline Ferralli, Yukio Nakamura, Sebastien Smallwood, Charisios D. Tsiairis

**Affiliations:** Friedrich Miescher Institute for Biomedical Research, Maulbeerstrasse 66, 4058 Basel, Switzerland; University of Basel, Petersplatz 1, 4001 Basel, Switzerland; SIB Swiss Institute of Bioinformatics, 4058 Basel, Switzerland; Institute of Medical Sciences, University of Aberdeen, AB25 2ZD Aberdeen, United Kingdom

## Abstract

Mechanical forces shape cell fate decisions during development and regeneration in many systems. Epithelial lumen volume changes, for example, generate mechanical forces that can be perceived by the surrounding tissue and integrated into cell fate decisions. Similar behavior occurs in regenerating Hydra tissue spheroids, where periodic osmotically driven inflation and deflation cycles generate mechanical stimuli in the form of tissue stretching. Using this model, we investigate how such mechanical input guides the de novo formation of differentiated body parts. We show that the expression of the organizer-defining factor Wnt3 functions as a quantitative readout of cellular stretching and, when supplied externally, enables successful regeneration without mechanical stimulation. This finding represents a previously undescribed cellular mechanism for converting mechanical stimuli to a biochemical signaling readout and guiding cell fate transitions. It also elucidates the role of mechanical oscillations in Hydra regeneration, which long remained unclear. The presence the Wnt/mechanics interplay in Hydra and its relatives underscores the ancient evolutionary history of this crosstalk, possibly extending back to the first metazoans. Since Wnt signaling crosstalks with cellular mechanics in various developmental and disease contexts, it can also represent a conserved feature of this signaling pathway.

---

Animal bodies and organs display an overwhelming variability of forms, yet their development relies on a relatively small repertoire of key processes such as folding, branching, and lumenization (Shahbazi, 2020; Teague et al., 2016). These morphogenetic processes often generate mechanical forces that can guide cellular behaviors (Hannezo and Heisenberg, 2019; Heisenberg and Bellaïche, 2013). For example, tissue stretching, resulting from lumen expansion, was shown to impact fate decisions of the stretched cells (Chan et al., 2019; Li et al., 2018; Ryan et al., 2019). To understand, how such mechanical cues crosstalk with biochemical signaling to drive cell identity changes, anatomically simple systems with well-defined cell differentiation processes, and amenable to experimental manipulation, are needed. Regenerating *Hydra* tissue is such a simple, experimentally tractable, system, where mechanics and cell fate decisions coincide but their connection has remained elusive (Vogg et al., 2019b).

When a small fragment is taken from the body of Hydra, the two-layered epithelial tissue folds into a spheroid with a lumen in the center. This spheroid gradually re-establishes the full body plan as the head (hypostome) with tentacles, and the foot (peduncle) of the animal appear, marking the opposite ends of its body axis (Fig. 1a). The uniformity of the epithelial cells is first broken when a small subpopulation starts expressing the Wnt3 ligand, differentiating into the head organizer which guides the morphogenesis of the surrounding tissue (Broun et al., 2005; Hobmayer et al., 2000). Interestingly, cycles of osmotically driven mechanical oscillations accompany the regeneration process (Kücken et al., 2008). Since *Hydra* is a freshwater animal, the cells need to maintain their osmotic balance against water that enters from the hypotonic environment. They excrete the surplus water into the spheroid lumen (Benos et al., 1977). Thus, the spheroid inflates, and the tissue is stretched at the same time. Once the stretching reaches a critical threshold, the tissue ruptures, causing the spheroid to deflate, and the cycle is repeated (Suppl. Video 1). In addition, the spheroid oscillatory behavior provides landmarks of regeneration progress, as it changes shortly after symmetry breaking and the mouth establishment (Soriano et al., 2009). The pressure accumulated during the inflation can be released through this newly formed weak spot, and the oscillation pattern transitions from a high-amplitude, low-frequency regime (Phase I) to a low-amplitude and high-frequency one (Phase II, Fig. 1c).

**Fig. 1.**
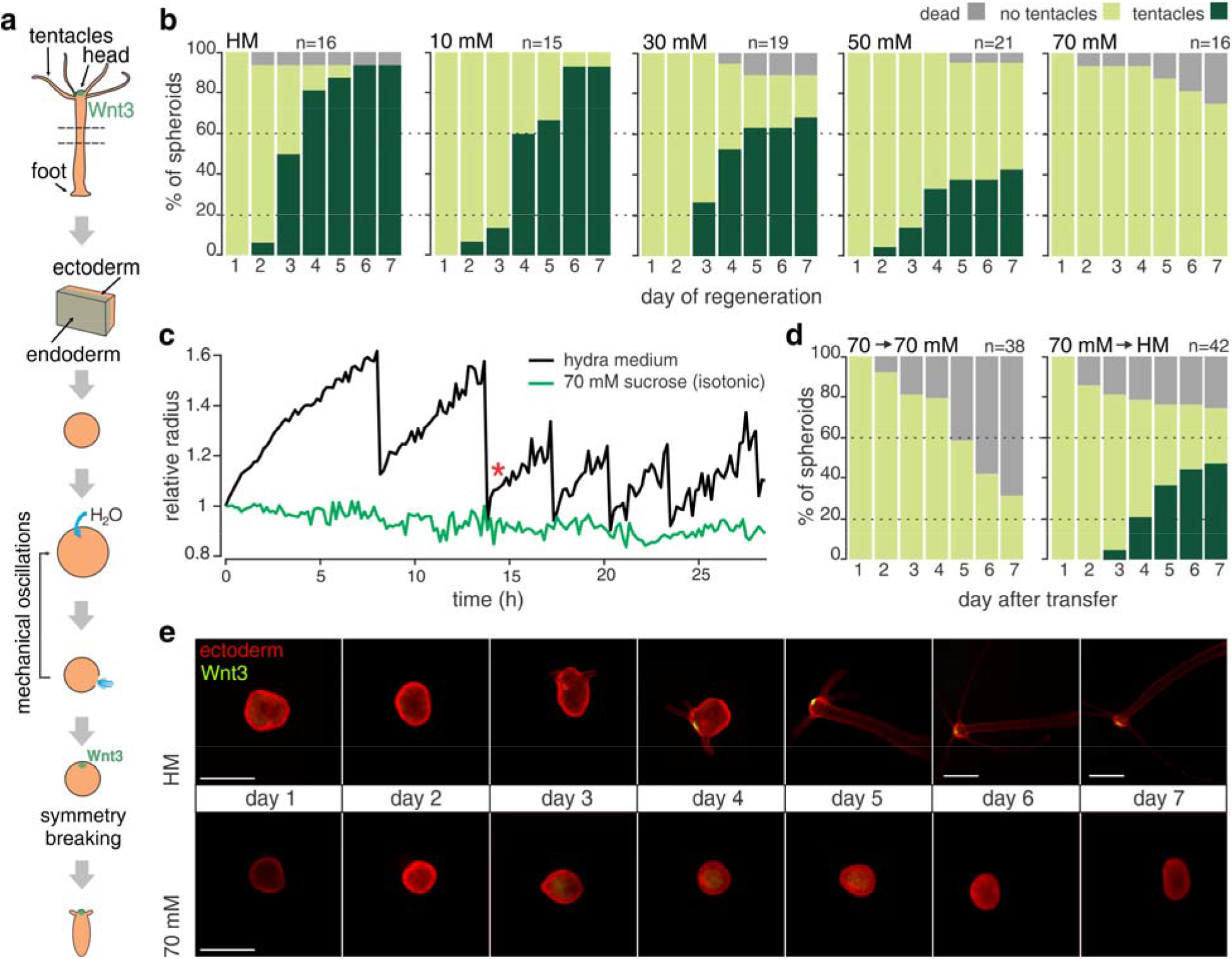
Mechanical stimulation is required for symmetry breaking and regeneration. (a) Spheroid preparation and regeneration. (b) Regeneration success decreases in media with increasing osmolarities. Cumulative plots for n animals from 3 independent experiments. The concentration of sucrose in hydra medium is indicated above each plot, 70 mM being isotonic. HM-hydra medium. (c) Quantifications of the size of representative spheroids from Movies S1-S2. Note the absence of oscillations in isotonic conditions. Asterisk marks PhaseI/II transition. (d) Regeneration resumes in HM after previous incubation in isotonic medium for 72 h. (e) Fluorescent imaging of representative spheroids from a Wnt3-reporter line in normal, and isotonic conditions without oscillations. There is a known delay in the reporter visibility, which causes it to be observable only after tentacle appearance(Nakamura et al., 2011). All scale bars, 500 μm.

Given the osmotic nature of inflation mechanism, oscillations can be perturbed by manipulating environmental osmotic pressure. Including additional osmolytes (e.g. sucrose) in the medium slows down the spheroid inflation rate in a concentration-dependent manner (Kücken et al., 2008), and in isotonic conditions, the oscillations cease completely (Suppl. Fig.1a, Suppl. Video 2). Using this approach, we observed a progressive loss of regeneration capacity (measured by the appearance of new tentacles) with increasing osmolarity. Under isotonic conditions (70 mM sucrose), spheroids completely fail to regenerate (Fig. 1b and Suppl. Fig. 1b). However, they remain viable for several days and can reinitiate the regeneration program when transferred back to normal medium (Fig. 1d), indicating that isotonic environment is not inherently toxic. Rather, it suggests that the absence of mechanical input puts a temporary halt to the regeneration program.

To answer whether mechanical input is required for the emergence of the Wnt3^+^ organizer cells, we used a Wnt3-GFP reporter line. Remarkably, all spheroids lacked GFP-positive foci under isotonic conditions (Fig. 1e, and Suppl. Fig. 1c). Thus, mechanical input is required for the differentiation of the head organizer rather than the execution of the downstream morphogenetic program of tentacle development. Surprisingly, even in the absence of oscillations, many spheroids elongate in the isotonic medium over time (Suppl. Fig.1d). Acquiring an asymmetric shape is, therefore, not a hallmark of symmetry breaking, as previously suggested (Soriano et al., 2009). Head regeneration in bisected animals is also hindered in isotonic conditions, indicating that some level of mechanical stimulation is a general requirement for *Hydra* regeneration (Suppl. Fig.1e).

To understand the role of these mechanical oscillations we examined the range of Phase I cycle numbers among spheroids. Such differences can result from intrinsic symmetry breaking variability among the tissue pieces. To investigate the source of this variability, we looked at the behavior of spheroids originating from different axial positions. Classical grafting experiments have demonstrated gradients of head formation and inhibition capacity along the axis. Tissue originating closer to the head is able to establish a new organizer more efficiently when grafted but, as a host environment, it is less conducive to a new organizer appearance (MacWilliams, 1983; Shimizu, 2012). Therefore, we examined whether the axial origin of the regenerating spheroids also impacts their mechanical dynamics. Indeed, spheroids derived from tissue closer to the head require more cycles before breaking the symmetry than those originating further away. These results thus likely reflect the persistence of an inhibitory property bestowed by the proximity of an existing head (Fig. 2a-b). In contrast, mechanical properties do not seem to be graded along the main axis, since the key oscillation parameters like amplitude and period appear similar for all the measured positions (Suppl. Fig.2a-d).

**Fig. 2.**
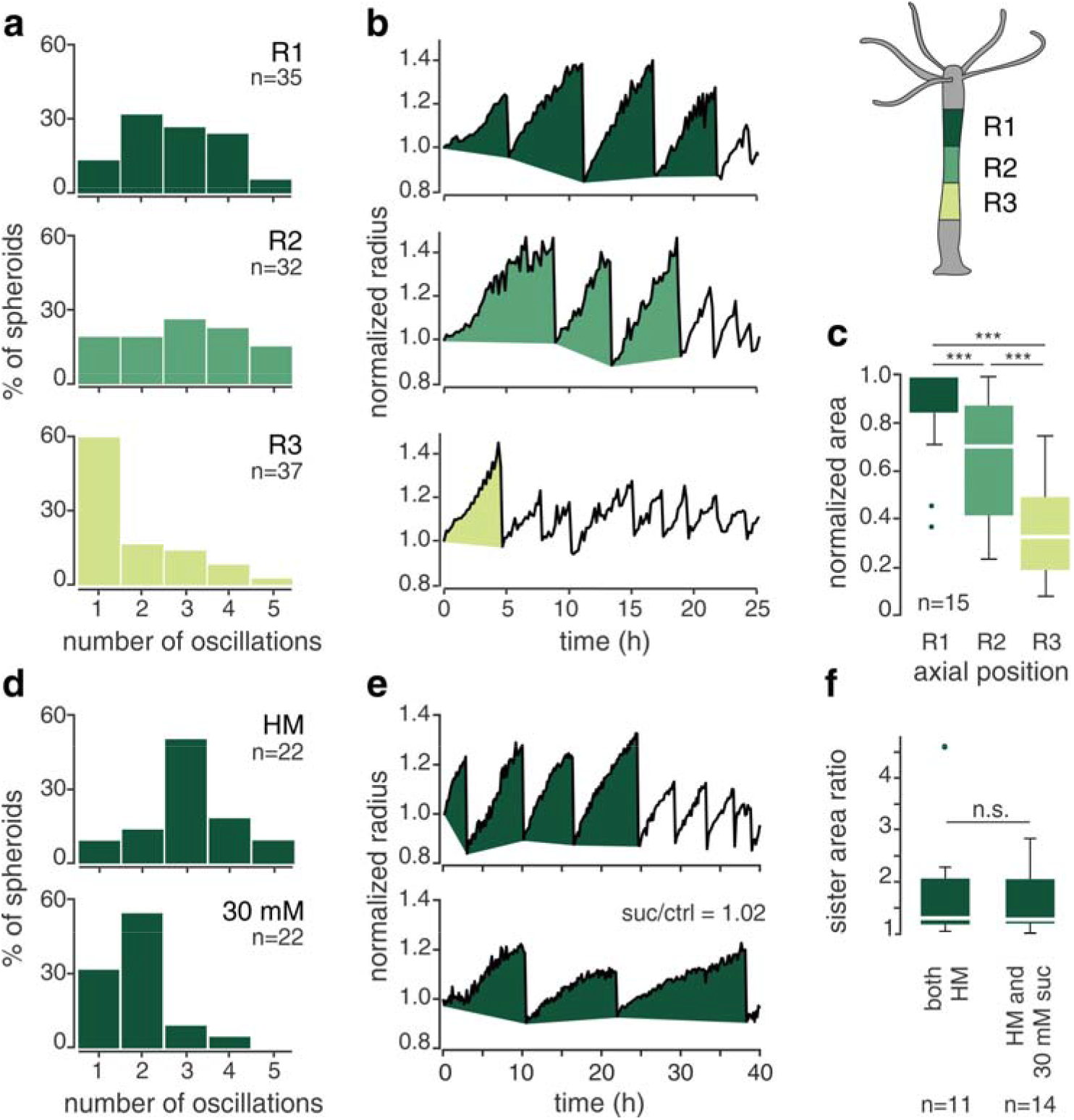
Spheroids integrate the amount of stretching over time. (a) Distributions of the number of Phase I oscillations for spheroids from different axial positions (R1 – R3, see schematic). Data of n samples pooled from 3 independent replicates. (b) Representative examples of oscillation patterns for three spheroids derived from different axial positions of the same individual. Filled parts correspond to Phase I oscillations. (c) Quantification of the Phase I area for 15 animals from (a) for which data were available for all three positions. Measurements for each animal were normalized to the R1 position. (d) Comparison of the oscillation number distribution for R1 spheroids in HM vs. 30 mM sucrose (n samples from 5 independent experiments). (e) A representative example of two spheroids from the identical axial position of a single animal (sister spheroids) showing a similar Phase I area in different media. (f) Area ratios for pairs of sister spheroids. They were either both cultured in hydra medium or one of the pair was put into 30 mM sucrose. The ratios for each pair are always normalized to the sister with a smaller Phase I area, irrespective of the conditions (n samples pooled from 3 independent experiments). Data in C and F were analyzed using the Mood’s median test. *** p < 0.0005 (p_R1/R2_=1.0322e-4, p_R1/R3_=2.9061e-08, p_R2/R3_=8.7409e-04)

Having established that mechanical input is indispensable for successful regeneration and that the number of Phase I oscillations correlates with an inhibitory axial gradient, we then asked whether spheroids require a specific number of cycles to be able to regenerate (a “counting” model). Alternatively, they might sense the overall amount tissue stretching experienced during the oscillations, disregarding deflation events (a “continuous” model). To distinguish between these options, we imaged spheroids in a medium with intermediate osmolyte concentration (30 mM sucrose), which slows down the rate of inflation and, consequently, prolongs each cycle’s duration (Suppl. Fig. 2e-h). Only spheroids from the R1 position were used, since they have the highest number of cycles on average. Assuming a counting mechanism, all cycles should be executed irrespective of their duration. Instead, we observed a decrease in the average number of Phase I oscillations in higher osmolarity (Fig. 2d). This argues that a continuous mechanism relying on the overall amount of tissue stretching seems more plausible. This metric can be approximated by quantifying the area under the plot of Phase I oscillations (hereafter Phase I area, Fig. 2b-c). Indeed, sister spheroids, derived from the same axial position and from the same animal, tend to have a very similar Phase I area even in different medium osmolarities (Fig. 2e-f). These results underscore the crucial role of sustained mechanical tissue stretching during inflation for successful symmetry breaking and regeneration.

To understand the molecular impact of tissue stretching, we combined a time course of RNA-sequencing in spheroids (Suppl. Fig. 3) with a positional RNA-sequencing in isolated body pieces from individual animals (Suppl. Fig. 4). The body parts’ transcriptomes allowed us to map gene expression profiles onto positional identities along the axis (Fig. 3a). When projected into this map, the time course data of >150 individual spheroids per condition revealed a difference in developmental trajectories. Under normal conditions, spheroids follow a relatively straightforward path from a body-like state towards a regenerated animal (Fig. 3b). However, in isotonic medium, without mechanical input, the trajectory lacks directionality and follows an undulating path that eventually approaches a foot-like identity (Fig. 3c). Remarkably, when examined for the characteristic foot peroxidase staining, about 50 % of the spheroids exposed to isotonic conditions are indeed positive for this foot marker (Suppl. Fig. 5). However, these spheroids will never accomplish head regeneration. The establishment of the foot thus seems largely independent from mechanical stimulation and head regeneration, supporting the view that the two poles of the oral/aboral axis might function as separate organizing centers (Meinhardt, 1993).

**Fig. 3.**
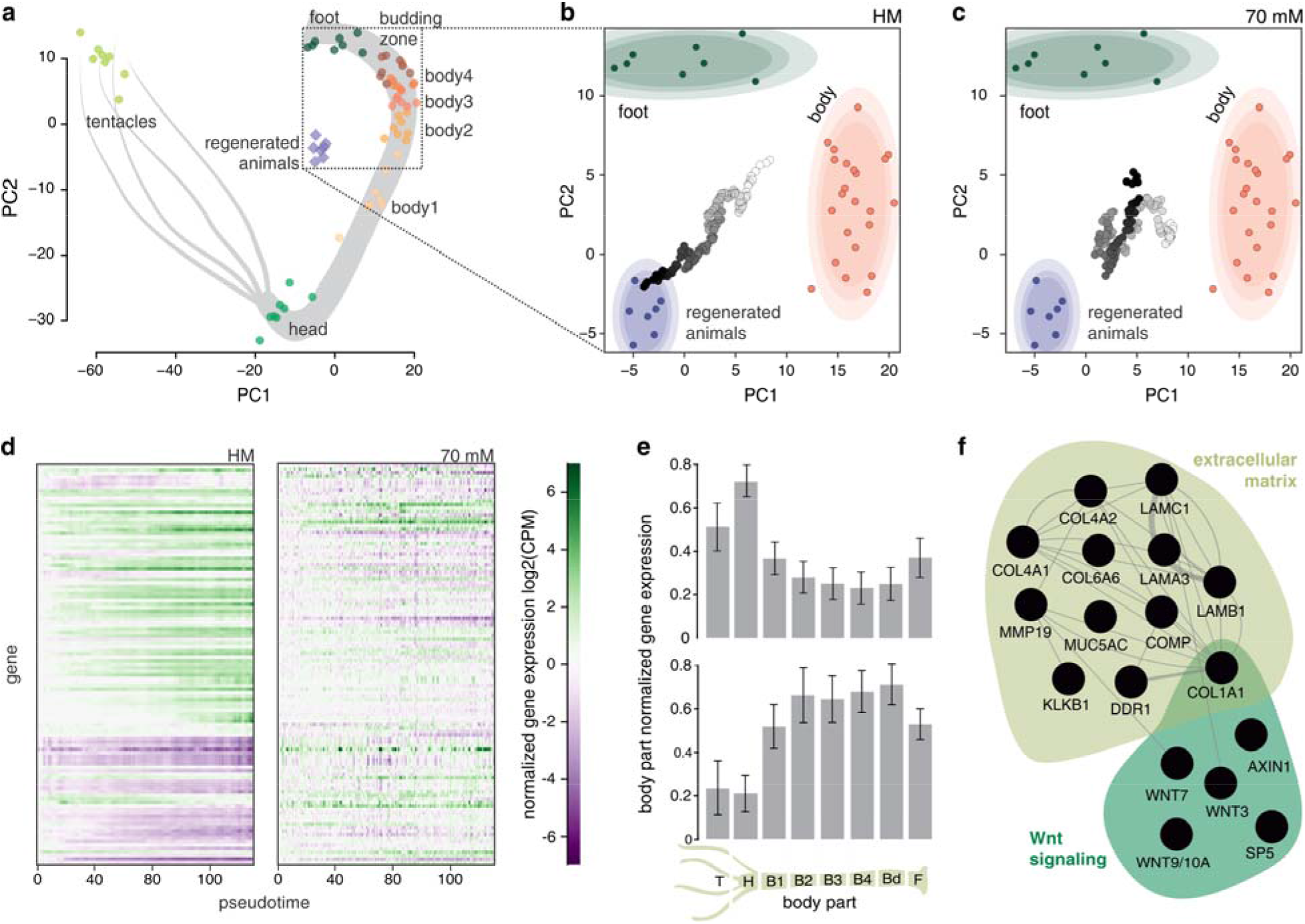
Head regeneration transcriptional program shows a strong mechanical dependence. (a) PCA plot of different body parts based on their transcriptional identity. Full animals regenerated from spheroids (harvested 72 h after cutting) are also included. Dotted box indicates the area enlarged in the following panels. (B – C) Projections of the spheroid time-course data into the body PCA space, with (b) and without mechanical input (c). Shaded ovals around the body parts (body2 – body4) and regenerated animals data correspond to 90 %, 80 %, and 70 % gaussian confidence intervals (light to dark shading). The color of the projected time-course data represents the pseudotime progression (from white to black). Each individual dot corresponds to a single spheroid. (d) Heatmaps of the behavior of the top 10 % (n=113) genes affected by lack of mechanical simulation in control and isotonic conditions. For each gene, the initial log2(CPM) value was subtracted. (e) Average expression patterns in the body for the increasing (upper panel) and decreasing genes (lower panel) from D. The expression data for each gene were maximum normalized to the body part with the highest expression of that gene, before averaging per body part. Error bars show 95 % confidence intervals. (f) Subnetworks of genes showing functional enrichment within the upregulated genes, affected by lack of oscillations. Edges indicate physical, genetic, and predicted interactions or coexpression in human cells.

We then looked at the top 10 % of genes (n=115) most severely affected by the removal of mechanical input (Suppl. Fig. 3d), to gain insight into the transcriptional changes behind the observed developmental differences. The majority (n=73) of these genes were increasing over time during normal regeneration, while a smaller portion (n=37) showed a decreasing trend (Fig. 3d). Interestingly, these groups of genes also have very different axial expression profiles. The rising genes show a clear enrichment in the head region, consistent with their role in setting up the body part most affected by the lack of mechanical oscillations. On the other hand, the group of decreasing genes shows enrichment in the undifferentiated body column, highlighting the failure to differentiate properly when oscillations are inhibited (Fig. 3e). Looking at functional annotations within the rising cluster (Suppl. Table1 and 2), we found an enrichment of extracellular matrix related genes, which likely reflects the tissue remodeling that has to occur in order to tolerate the mechanical stress and to regenerate. More importantly, several components of the Wnt signaling pathway were also enriched, including three ligands of the canonical Wnt signaling (Fig. 3f).

Further investigating the impact on Wnt signaling, we noticed that the lack of mechanical stimulation affected the expression dynamics of all the canonical Wnt signaling ligands (Fig. 4a, Suppl. Fig. 6a). These genes are activated sequentially during normal regeneration in bisected animals (Lengfeld et al., 2009) and we observed a similar progression for spheroids. After the pseudotime point 80 most of the transcripts rapidly increase, reflecting a successful establishment of the mouth organizer. However, none of these changes takes place without mechanical stimulation. Interestingly, Wnt3 expression has a behavior distinct from the other Wnt ligands (Fig. 4b). Previous studies have shown that Wnt3 is upregulated very early in response to injury (Chera et al., 2009; Vogg et al., 2019a) and then sustained throughout the regeneration process. Yet, in isotonic conditions, its expression fails to be sustained and rapidly decreases. We therefore propose that the Wnt3 transcriptional output functions as a readout of the mechanical input. We tested this hypothesis by performing a qPCR time course under different osmolarities, corresponding to different intensities of mechanical stimulation. As expected, the ability to sustain Wnt3 expression was anticorrelated with the mechanical input strength (Fig. 4c, Suppl. Fig. 6b-c).

**Fig. 4.**
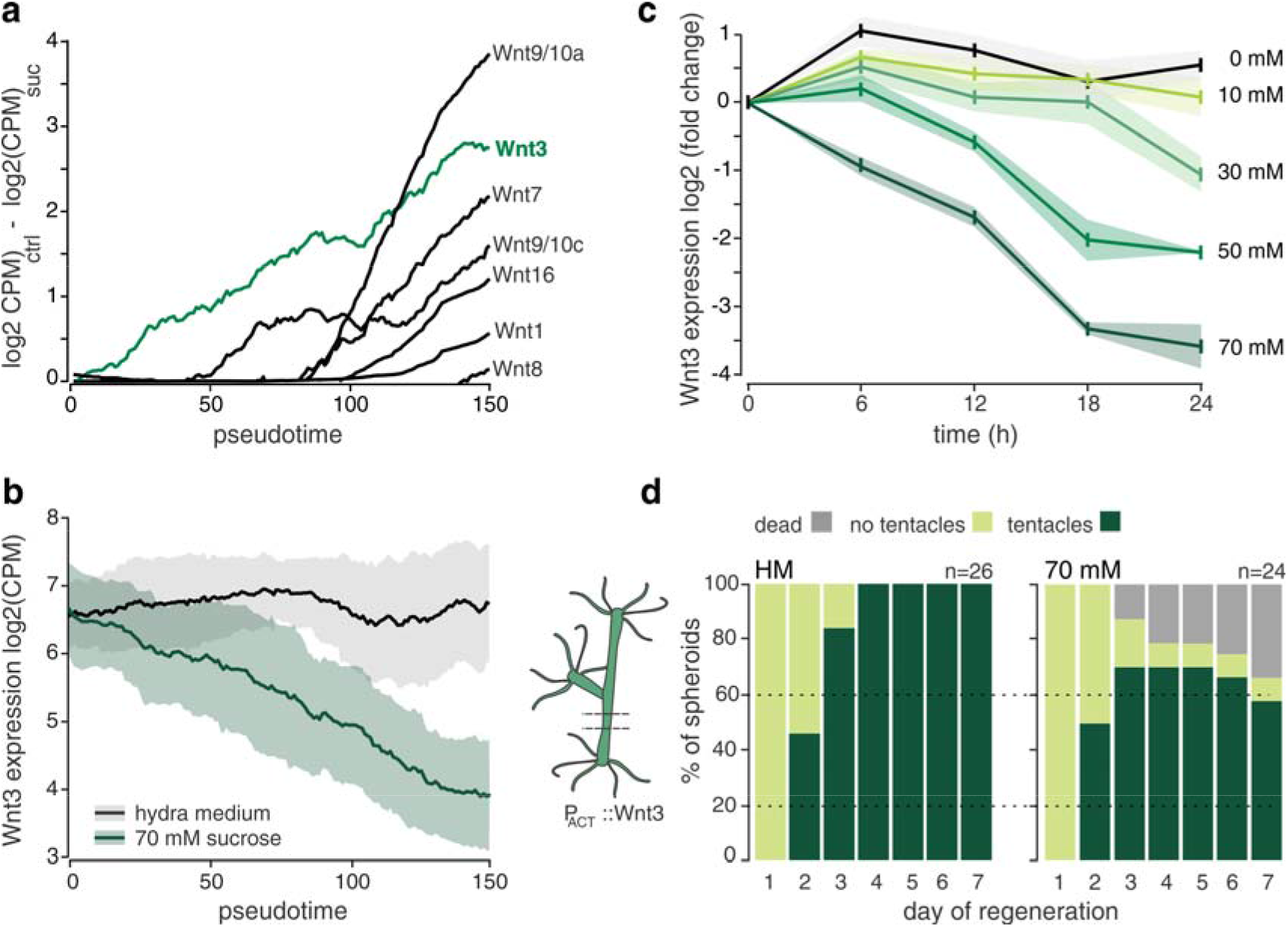
Wnt3 expression is a quantitative readout of mechanical stimulation. (a) Temporal development of the expression differences of canonical Wnt ligands between control and isotonic conditions. (b) Wnt3 expression patterns in HM and isotonic medium. In both A and B the lines indicate moving averages (moving window width is 36 pseudotime points). Shaded area represents a moving standard deviation (window width identical to the mean). (c) Wnt3 expression dynamics in different osmolarities, measured with qPCR. Data for each condition were normalized to the initial time point. Shown are averages from 3 independent biological replicates. Shaded areas represent standard error of the mean. (d) Rescue of regeneration without mechanical stimulation in Wnt3-overexpressing spheroids. Cumulative plots from 3 independent experiments.

We then asked whether providing Wnt3 would be sufficient to rescue regeneration in the absence of mechanical stimulation. Indeed, both spheroids derived from Wnt3 overexpressing animals (Suppl. Fig. 7), as well as those derived from early buds that already possess an organizer, were able to regenerate in isotonic conditions (Fig. 4d, Suppl. Fig. 6d-f, Suppl. Video 3 – 6). We thus conclude that sustaining Wnt3 expression is crucial for successful *Hydra* head regeneration and that it is made possible by stretching-dependent transcriptional activation of its gene.

Our work provides insights into the role of mechanical input in Hydra regeneration and its connection to the underlying molecular pathway, which remained elusive despite the fact that the oscillations are a prominent feature of the process (Braun and Keren, 2018; Ruiz-Herrero et al., 2017). This opens several exciting avenues for future research, such as investigating how is the tissue stretching converted to Wnt3 transcriptional changes, or how is the organizer spatially specified. Moreover, since fluid-filled lumens are one of the leitmotifs of animal morphogenesis (Ruiz-Herrero et al., 2017), our system offers an excellent platform to study the general rules of lumen formation and function, in particular the relationship between pressure-dependent cell stretching and cell fate decisions (Hannezo and Heisenberg, 2019). While mechanisms similar to the one described here could work with many signaling molecules, the case of Wnt is intriguing thanks to its ancient evolutionary past. The origin of Wnt signaling coincides with the origin of multicellularity (Loh et al., 2016), thus suggesting that its mechanosensitivity in Hydra and other cnidarians (Pukhlyakova et al., 2018) reflects an early evolutionary connection. Whereas first multicellular animals likely had a poor signaling repertoire, the combination of biochemical signals with mechanical cues may have enabled body plan elaboration (Newman et al., 2006), as we report here for Hydra spheroids. Importantly, mechanical input driving Wnt expression is a widespread phenomenon in diverse developmental contexts (Baron and Kneissel, 2013; Cha et al., 2016; Shinozuka et al., 2019) as well as in disease (Fernández-Sánchez et al., 2015). We thus suggest that this might be a conserved mode of Wnt signaling activation through mechanical stimulation.

## Acknowledgments

We thank the members of the FMI Functional Genomics Facility for performing the RNA-sequencing, the members of the Facility for Advanced Imaging and Microscopy for their support to the imaging part of this work, and Thomas Bosch (Christian-Albrechts-Universität, Kiel) for providing the AEP ecto pAct::GFP *Hydra* line. We are also thankful to our colleagues Prisca Liberali, Susan Gasser, and Fabian Braukmann for their constructive comments that helped us improve the manuscript. This work was supported by the Novartis Research Foundation.

## Author contributions

JFc and CDT designed the study and wrote the manuscript. JFc, JFi, YN, SS, and CDT perfomed the experiments. JFc, PP, and CDT analyzed the data.

## Methods

### Animal strains and culture conditions

All experiments were performed using either the AEP of 105 strains of *Hydra vulgaris.* Animals were kept in hydra medium (HM, 1mM CaCl_2_, 0.2 mM NaHCO_3_, 0.02 mM KCl, 0.02 mM MgCl_2_, 0.2 mM Tris-HCl pH 7.4) at 18 °C and fed with freshly hatched *Artemia* nauplii 3 times a week. Individuals without buds, starved for 24 hours, were used for experiments, if not indicated otherwise.

### Spheroid preparation

Details of spheroid generation, imaging and oscillation analysis can be found in a previously published protocol (Ferenc and Tsiairis, 2020). Animals were bisected (exact position of the initial cut for different experimental designs is detailed below) and one or several thin tissue rings were obtained by sequential cutting. The rings were then split in 2 – 3 rectangular pieces that were left to close for ca. 2 – 2.5 h in HM or dissociation medium (DM, 3.6 mM KCl, 6 mM CaCl_2_, 1.2 mM MgSO_4_, 6 mM sodium citrate, 6 mM sodium pyruvate, 4 mM glucose, 12.5 mM TES, pH 6.9) at room temperature. Properly closed spheroids (typical diameter 300 – 500 μm) were then separated from the pool and used for further experiments.

### Live imaging of developing spheroids

Multi-chamber LabTek slides (Nunc) were used for imaging. Spheroids were placed in individual agarose wells (ca. 1 mm in diameter) within the chambers, to retain them in the field of view. The wells were created by coating the bottom of the chambers with a 2-3 mm thick layer of 1% agarose in HM and puncturing holes using a 1 ml micropipette tip after the gel solidified. The chamber was then filled with the desired imaging medium. In case of sucrose treatments, the agarose gel contained sucrose in concentrations equal to the overlaying medium. For inhibitor treatments, drug concentration in the imaging medium was adjusted, taking the volume of agarose into account. To prevent diluting the imaging medium in the chamber, closed spheroids were first transferred to a dish with bigger volume (ca. 10 ml) of the imaging medium and positioned in the imaging wells only after this wash. Samples were imaged every 10 min at 21 °C under brightfield illumination using the Nikon Ti2-Eclipse microscope with a 10x CFI Plan Apochromat Lambda objective (Nikon) and iXon-Ultra-888 EMCCD camera (Andor). Images were acquired in the 1024×1024 format, with pixel size of 1.3 μm.

### Quantification of oscillation parameters

Brightfield images were batch segmented in Fiji (Schindelin et al., 2012) A course median filter (radius = 60 px) was first applied, followed by Phansalkar local thresholding (radius = 50 px, r and k parameters = 0) and another round of median filtering (radius = 20 px). The area of segmented objects bigger than 5000 μm^2^ including holes was then measured. Radius was calculated from these values, assuming a spherical object. Radius data for each sample were normalized to the starting value and further processed in Matlab using custom functions^31^ to extract the slope, amplitude and period of the Phase I oscillations. The number of oscillations and Phase I/II transition was determined manually by two independent experimenters.

### Scoring regeneration and Wnt-center appearance in spheroids

Spheroids from the lines *AEP ectop Act::eGFP* (Wittlieb et al., 2006) and *AEP ectop Act::dsRed, pWnt3::eGFP* (Nakamura et al., 2011) were prepared as described above. Animals were bisected at 20% of the body length below the head and one or two rings per animal were generated. Spheroids were left to close in HM. Similar to previous studies (Kücken et al., 2008), we used sucrose to alter the medium osmolarity. Sucrose was dissolved in HM to the desired concentration (0, 10, 20, 30, 50 and 70 mM (isotonic)), supplemented with antibiotics (50μg/ml kanamycin, 100μg/ml streptomycin) and filter sterilized. Spheroids were randomly split among the conditions (0 and 70 mM only for Wnt3-reporter line) in 24-well plates, kept at 18 °C, and scored for the presence of tentacles in 24 h intervals for 7 days. Additionally, the presence of the organizer was assessed in the Wnt3-reporter line spheroids by fluorescence imaging. The Zeiss AxioZoom V.16 fluorescent stereomicroscope with the Zeiss Axiocam MRm CCD camera was used. Finally, the regenerated animals/surviving spheroids were fixed on the 7th day of the time course and stained to determine the presence of the foot.

### Experiments with restarted oscillations

*AEP ectop Act::eGFP* spheroids were prepared as described above, and the whole population of properly closed spheroids was transferred to 70 mM sucrose in HM to prevent mechanical oscillations. Samples were kept in this medium for 72 h at 18 °C. Surviving spheroids were then randomly split into two groups and either transferred to 70 mM sucrose again (controls) or to HM, allowing the mechanical oscillations to restart. Both populations were monitored daily for tentacle appearance as in the previous experiments.

### Scoring head regeneration in bisected animals

*AEP ectop Act::eGFP* animals were bisected at 50% body length and the halves were kept separately in the wells of 24-well plates at 18 °C. Tentacle appearance was monitored in 24 h intervals in the foot halves for 5 days.

### Peroxidase foot staining

The staining was performed as previously described (Hoffmeister and Schaller, 1985). Briefly, animals were relaxed in 2 % urethane in HM and fixed with 4 % paraformaldehyde in HM at 4 °C overnight. Animals were then washed in PBS + 0.1 % Tween-20 for 5 min and stained for 15 min in the staining solution (0.02 % diaminobenzidine, 0.03 % hydrogen peroxide, 0.25 % Triton-X in PBS). To stop the enzymatic reaction, samples were washed once again in PBS+Tween for 15 min. The whole staining procedure was performed at room temperature with mild agitation.

### Imaging and quantification of oscillations in spheroids of different axial origin

To obtain spheroids from different axial positions, the head of the *AEP ectop Act::GFP* animals was cut away at ~ 10 % of the body length and three rings of tissue were obtained sequentially. Each ring was split into two pieces. Pieces were incubated at room temperature for 2.5 h in DM while closing. To allow backtracking of the piece identity, sister pieces from each ring were kept in separate wells of a 24-well plate, noting the axial position and animal of origin. Similar arrangement was followed for imaging, which otherwise proceeded as described above. Quantifications of oscillation parameters from obtained time-lapse images were also performed as outlined previously.

### RNA-sequencing time course of developing spheroids

Over the course of 10 h, few dozens of spheroids were prepared every hour from the 105 strain, let to close for 2 h in DM and then transferred to either HM or HM with 70 mM sucrose. 8 spheroids from each batch were then collected 3 h and 13 h after cutting the last batch, thus creating a time course spanning a window of 22 h. Spheroids were collected individually in 96 well plates in 45 μl of RLT+ buffer (Qiagen) and stored at −80 °C for a later RNA purification. This was performed using the Zymo Direct-zol MagBead reagents following a modified manufacturer protocol. Briefly, 45μl of Zymo Binding Buffer and 4μl of MagBeads were added to the samples/RLT+ buffer and incubated for 10min. Beads were washed 3 times with 100% ethanol and incubated with 12.5μl of DNAseI (Zymo) for 10min. The RNA was captured back on the beads using 100μl of MagBead PreWash buffer (10min incubation). After washing the beads 3 times with 100% ethanol, elution was performed with water (13μl) at 55C for 15min. All above incubations were performed at room temperature on a plate shaker unless specified. The resulting RNA quality was assessed using bioanalyser or tapestation RNA HS kit, and concentration determined using Qbit. cDNA amplification was performed using the SmartSeq2 approach as per the original protocol (Picelli et al., 2014b). Full length cDNA was processed for Illumina sequencing using Tagmentation with an in-house purified Tn5 transposase (Picelli et al., 2014a): 1ng of amplified cDNA was tagmented in TAPS-DMF buffer (10mM TAPS pH8.5, 5mM MgCl2, 10% DMF), at 55C for 7min. Tn5 was then stripped using SDS (0.04% final) and tagmented DNA was amplified using Phusion High-Fidelity DNA Polymerase (ThermoFisher). PCR was performed in the Phusion HF buffer, with a first extension at 72 °C for 3min, followed by 10 cycles of amplification (95 °C −30 s, 55 °C −30 s, 72 °C – 30s). Commercial Nextera XT indexes were used for the PCR amplification (1/5 dilution). Final libraries were sequenced on a HiSeq2500 (50cycles single-end) and demultiplexing performed using a standard Illumina bcl2fastq2 pipeline.

### Positional RNA-sequencing

Animals of the 105 strain were first bisected at the midpoint between the head and the budding zone. The halves created this way were bisected again in the middle between the head and the previous cut, or the previous cut and the budding zone, respectively, thus creating “body 2” and “body3” segments. Head with tentacles and budding zone with the foot were then removed from the remaining pieces to generate “body1” and “body4” segments. Finally, the budding zone was separated from the foot, using the change of endoderm coloration as a guideline for sectioning. Tentacles were also separated from the head, trying to remove as much of the tentacle tissue as possible. The individual segments were lysed immediately after being cut in 350 ul of RL buffer (Single Cell RNA Purification Kit, Norgen), supplemented with 1 % β-mercaptoethanol, frozen on dry ice and stored at −80 °C for later RNA isolation. Tentacles of a single animal were pooled as one sample. RNA extraction was performed according to the manufacturer’s instructions. Downstream processing of the isolated RNA was identical with the previously described experiment, using the SmartSeq2 protocol and in-house Tn5 transposase. RNA from animals regenerated from spheroids was also isolated and sequences as described here. The spheroids were prepared from the 105 strain, as described in the time course sequencing experiment above and left to regenerate in HM for 72 hours. Whole regenerated animals were then collected in 350 ul of RL buffer in three independent replicates (2-3 animals/replicate).

### RNA-seq data alignment and pre-processing

The SmartSeq2 libraries for the spheroid RNAseq generated a total of ~8.9 billion 50nt long reads (accounting for an average sequencing depth of ~25 million reads per spheroid). The Illumina Smartseq2 libraries for the positional and regenerated animals RNA sequencing generated a total of 1.7 billion and 250 million 50nt long reads respectively corresponding to an average depth of ~27 million reads per segment replicate and ~31million reads per regenerated animal. After demultiplexing, reads were aligned against the hydra genome guided by transcriptome annotation (NCBI Hydra vulgaris assembly Hydra_RP_1.0, NCBI Hydra vulgaris annotation release 102) using STAR *(7)* version 2.5.0 with command line parameters: — *outSJfilterReads Unique --outFilterType BySJout --outFilterMultimapNmax 5 -- alignSJoverhangMin 8 --alignSJDBoverhangMin 4 --outFilterMismatchNoverLmax 0.1 -- alignIntronMin 20 --alignIntronMax 1000000 --outFilterIntronMotifs RemoveNoncanonicalUnannotated --seedSearchStartLmax 50 --twopassMode Basic -- genomeChrBinNbits 12 --genomeSAsparseD 2 --quantMode GeneCounts.* Samples with library sizes smaller than 5 million reads were discarded from all subsequent analyses. In total 32/352 spheroid libraries and 1/72 animal segment libraries were removed at this step. The produced gene count tables for all remaining samples were library normalized after excluding from the size factor calculation the top 5 percentiles of highly expressed genes. As the samples along the 22h timeme-course of the spheroids were obtained in two collection sessions (one session for timepoints 1h-11h and one session for timepoints 12-22h), a batch effect was introduced. This manifested as a slight discontinuity both in PCA projections of the samples and in select timecourse expression profiles of individual genes. In order to mitigate this effect, we took advantage of the fact that spheroids develop asynchronously (median pseudotime spread per collection point of 1.6h, see section on pseudotime ordering) allowing us to pinpoint samples across batches that nearly overlap in terms of dynamics. The correlation (Spearman’s rho) of samples from adjacent timepoints across the two collection series (16 samples collected at 10-11h and 14 samples collected at 12-13h) was calculated to identify pairs of top-3 mutual nearest neighbors (MNNs), corresponding to inferred overlapping samples across batches. These MNN pairs were used to calculate average gene-specific shifts that were then applied as correction factors on all samples from the two batches. This simple strategy efficiently removed the observed discontinuities both in sample projections and individual gene profiles. The corrected spheroid expression values are used for all downstream analyses.

### Pseudotime ordering of spheroids

Individual spheroids evolve asynchronously, with samples collected at a specific experimental time exhibiting technical but also internal developmental time variation. In order to recover the underlying gene expression dynamics, it is therefore necessary to order the samples along an axis corresponding to the temporal evolution of the system and to obtain latent coordinates for each sample on that axis. Our strategy for pseudotime ordering relied on first identifying data features that are smooth functionals of the time variable and subsequently using those features to obtain a 1d projection of the spheroids on a basis that corresponds to the pseudotime axis. This procedure was applied separately to the regular medium and isotonic 22h spheroid time-course datasets sampled at 1h intervals. We begin by selecting the top 50% most variable genes, according to a within-dataset mean-variance trend fit. Each of the first 50 principal components (*P*) of this filtered dataset were fit against a generalized additive model using time (*t*) as the independent variable (function *gam* from the *mgcv* CRAN library, call: *gam(P ~ s(t), method = “GCV.Cp”, gamma=1.0)*). The dataset is then reconstructed using only components with gam-fit fdr-adjusted p-values < 0.05 as the rest will contain information almost orthogonal to time dynamics. Next we iterated over all 16, overlapping, 7h time windows and repeated the gam-fitting procedure for the expression of all genes (g) to identify individual features that are informative for time-ordering within the corresponding timeframe (function call: *gam(G ~ s(t, k=4), method = “GCV.Cp”, gamma=1.25)*). For each 7h time window, 100 1d isomap embeddings were performed using random subsets of half of the respective selected gene expression profiles. Within each 7h window, pseudotime was estimated as the mean of the returned 1d coordinates from the isomap iterations scaled to 7h plus a shift corresponding to the first window timepoint. This procedure returns between 1 and 7 pseudotime estimates per spheroid (depending on how many 7h windows cover the corresponding collection time). A final consensus pseudotime was obtained using the weighted average of the individual estimates using weights determined by the number and gam-fit significance of the genes selected in each respective 7h window.

### Projection of segment, regenerated animal, and spheroid data on a common subspace

For the common subspace projection of spheroid, segment and regenerated animal data expression values were first log2 transformed after smoothing using a pseudocount of 8 to shrink effect sizes of lowly expressed genes. The pseudotime-ordered spheroid data were further smoothed using a moving average with a window size of 16 (corresponding to a real-time window of ~2h) since spheroids exhibit considerable technical variation. We then obtained a single eigenvector basis by applying principal component analysis on the animal segment data (base R function *prcomp* with parameters *center=TRUE, scale=FALSE).* Only the intersection of the genes with at least a 2-fold change in gene expression (max absolute delta of 1 for the log transformed values) in both the segment and spheroid datasets were used. Finally, the spheroid and regenerated animal data were projected back to the eigenvector space determined by the animal segment data.

### Differential analysis of gene expression in spheroids in hypotonic and isotonic media conditions

In order to identify genes that are differentially expressed in the hypotonic (HM) and isotonic (70mM glucose medium) conditions we fitted a generalized additive model (GAM) on the delta of the two pseudo-time ordered (PTO) expression time series. First, in order to allow comparisons between the two conditions we had to account for the fact that, after filtering low-depth libraries, we end up with a larger number of spheroids in HM conditions compared to the 70mM conditions (170 vs 150 spheres, see section on RNA-seq data pre-processing). We downsampled the HM data by interpolating to 150 sampling points (*t*) to acquire two equal-length time series. The difference *Δ* of the two signals for every gene was scaled and fitted to a GAM (function *gam* from the *mgcv* CRAN library, call: *gam(Δ ~ s (t, k=8), method = “GCV.Cp”, gamma=1.0)*). In addition, we calculated effect sizes for each gene between the two conditions as the sum of the absolute values of the difference between the two mean-normalized signals, excluding the highly variable first three timepoints. This procedure allows us to identify genes that differ either in terms of scale or in terms of shape in their gene expression dynamics. Both the fdr-adjusted p-values from the fit (<1e-6) and the calculated effect sizes (>40) were used to select for genes differentially expressed in the two conditions.

### Analysis of the changing genes sensitive to mechanical stimulation

The 2269 genes that change their expression at least 2-fold during normal regeneration were clustered in 5 clusters using the *kmeans* function in Matlab. Before clustering, the gene expression time-series of each gene was mean-normalized. To select the top genes sensitive to the removal of mechanical stimulation, we only considered the ones with adjusted P-value <10^-6^. From this pool, the top 10 % were selected based on the sum of differences (n=113). To enable the functional analysis of these genes, their mammalian homologs were then annotated using the available data from *Hydra* genomics databases (see Table S1). GO term enrichment analysis was then performed using GeneMania (genemania.org) separately for members of clusters 3 and 5. Since there were only 3 genes that were members of other clusters, these were left out of the analysis.

### Real-time PCR time course of Wnt3 expression

Spheroids of the *105* strain were prepared as described previously, randomly split into 5 groups and incubated in media with different sucrose content (0, 10, 30, 50, 70 mM) at room temperature. Every 6 hours, a sample of 10 spheroids was randomly taken from each of the conditions, lysed in 350 ul of RL buffer with 1 % β-mercaptoethanol (Single Cell RNA Purification Kit, Norgen), frozen on dry ice and stored at −80 °C. After collecting all the samples, RNA was isolated as outlined in the instructions for the kit. The concentration and purity of the extracted RNA was verified using NanoDrop 1000 (Thermo Fisher Scientific). Next, cDNA was prepared with the Oligo(dT)12-18 Primer (Thermo Fisher Scientific) using the High-Capacity cDNA Reverse Transcription Kit (Applied Biosystems) as per manufacturer’s instructions. The qPCR reactions were carried out using the StepOnePlus Real-Time PCR cycler (Thermo Fisher Scientific) with a standard run method. Each reaction contained 10 ng of cDNA, 1x the Platinum SYBR Green qPCR SuperMix-UDG w/ROX (Invitrogen) master mix with 0.25 μM primers for either *hy-GAPDH* (fwd: 5’ – GACTTGGCCGTATTAACTTGAGC-3’, rev: 5’-CTACAAACAAGACGCCCTATTCG-3’) or *hy-WNT3* primers (fwd: 5’-TGCAGAAGGAATACGACTGGG-3’, rev: 5’-TGCTGGCTGTTGTAATAATTGGG-3’) in a total volume of 25 μl. The quantification of gene expression was performed using the StepOne software v 2.3 (Thermo Fisher Scientific).

### Generation of transgenic animals overexpressing Wnt3

The transgenesis of animals was performed as previously described (Nakamura et al., 2011; Wittlieb et al., 2006). Briefly, 2-cell stage embryos were injected with the pAct::GFP, pAct::Wnt3 construct and left to develop. After hatching, the polyps were reared separately and screened for the expression of the transgenesis marker GFP. Fully transgenic animals were then obtained from buds originating in the transgenic cell patches.

### Rescue experiments with buds and Wnt-overexpressing spheroids

Animals of the line *AEP ectop Act::eGFP* with buds in stages 3-4 (staging according to 37) were selected, and tissue rings containing the forming buds were cut. Only the budding fragment of the ring was then excised and allowed to form a spheroid in HM. Given the branched multiheaded morphology of the Wnt-overexpressing *AEP ectop Act::eGFP, pAct::Wnt3* line, it was not possible to cut tissue rings in a specific axial position. Rings were harvested from sufficiently long body fragments instead and cut into 2-3 fragments as usual. Fragments were left to close in HM. Both types of spheroids were then split between 0 mM and 70 mM sucrose in 24-well plates and scored for tentacle appearance, as detailed above.

### Whole mount in situ hybridization

In situ hybridization was performed according to previously published protocols (Bode et al., 2008) with minor modifications. Briefly, animals were fixed at 4 °C in 4% PFA overnight and then dehydrated in a series of 5 min washes in 25-50-75-100% methanol and incubated overnight in methanol at −20 °C. Rehydration then followed by successive 10 min washes of 75-50-25-0 % methanol in PBS + 0.1 % Tween 20 (PBT). Samples were then treated for 10 min with a 10 μg/ml Proteinase K solution and the reaction was stopped by incubation with 4 mg/ml glycine for 10 min. After 2x 5 min washes with PBT and 2x 5 min washes with 0.1 M triethanolamine solution, samples were treated with acetic anhydride for 2x 5 min, washed in PBT (2x 5min) and refixed in 4% PFA for 20 min. Thorough washing with PBT (5x 5 min) then followed and the animals were heat treated for 30 min at 80 °C. Following preincubation with the hybridization solution for 2 h at 55 °C, DIG-labelled RNA probe dissolved in hybridization solution was added. We used the same *Brachyury* probes previously described (Broun et al., 2005). The samples were incubated with the probes for 3 days and then washed with a series of 75-50-25-0 % solutions of hybridization buffer in 2x SSC. After blocking the samples in 20% sheep serum for 2 h at 4 °C, an incubation with the alkaline phosphatase conjugated anti-DIG antibody followed (overnight, 4 °C). To remove the unbound antibody, 8x 1 h washes with maleic acid buffer were performed, followed by an overnight wash in the same buffer. The next day, samples were treated by NTMT and levamisole, as detailed in the original protocol. Finally, the chromogenic reaction was performed using the BCIP/NBT Color Development Substrate (Promega) according to the manufacturer’s instructions.

### Transplantation experiments

Transplantations were performed on glass needles, hand-pulled from glass capillaries. Recipient wt AEP animals were bisected at 50% body length and the foot half was driven on the needle longitudinally. For controls, a ring of tissue from the same axial position was cut from the *AEP ecto pAct::eGFP* animals and threaded on the needle. Since it is not possible to match axial positions in the Wnt3 overexpressing animals, we used rings of tissue coming from anywhere in the branched body columns of these animals. The head half of the recipient was then added, thus sandwiching the transplant between the host body halves. Pieces were then secures with a piece of parafilm to prevent sliding of the needle and left for ~ 2h in HM to establish adhesion. After this time, healed grafts were carefully slid off the needle using fine forceps and left to recover. The success of transplantation was evaluated after 24h. Successful transplant were then assessed for ectopic head formation on day 5 after transplantation.

### Statistical analysis

All experiments have been performed in at least 3 independent replicates. Details about sample numbers are given in the figures or descriptions of the relevant experiments. Since the nature of the collected data often led to distributions that were not Gaussian, we used the nonparametric Mood’s median and Wilcoxon rank sum tests. All tests were performed as two-sided. Other relevant details are given in the figure legends and experiment descriptions. In all box plots, the central mark corresponds to the median, and the bottom and top box edges indicate the 25^th^ and 75^th^ percentiles, respectively. Whiskers show the range of the data points not considered outliers. Outliers are defined as points that have a value of less than *q_1_* – *w* × (*q*_3_ – *q*_1_), or more than *q_3_* + *w* × (*q*_3_ – *q*_1_). Here, *w* is the maximum length of the whisker, and *q*_1_ and *q*_3_ correspond to the 25th and 75th percentiles, respectively.

---

**Suppl. Fig.1.**
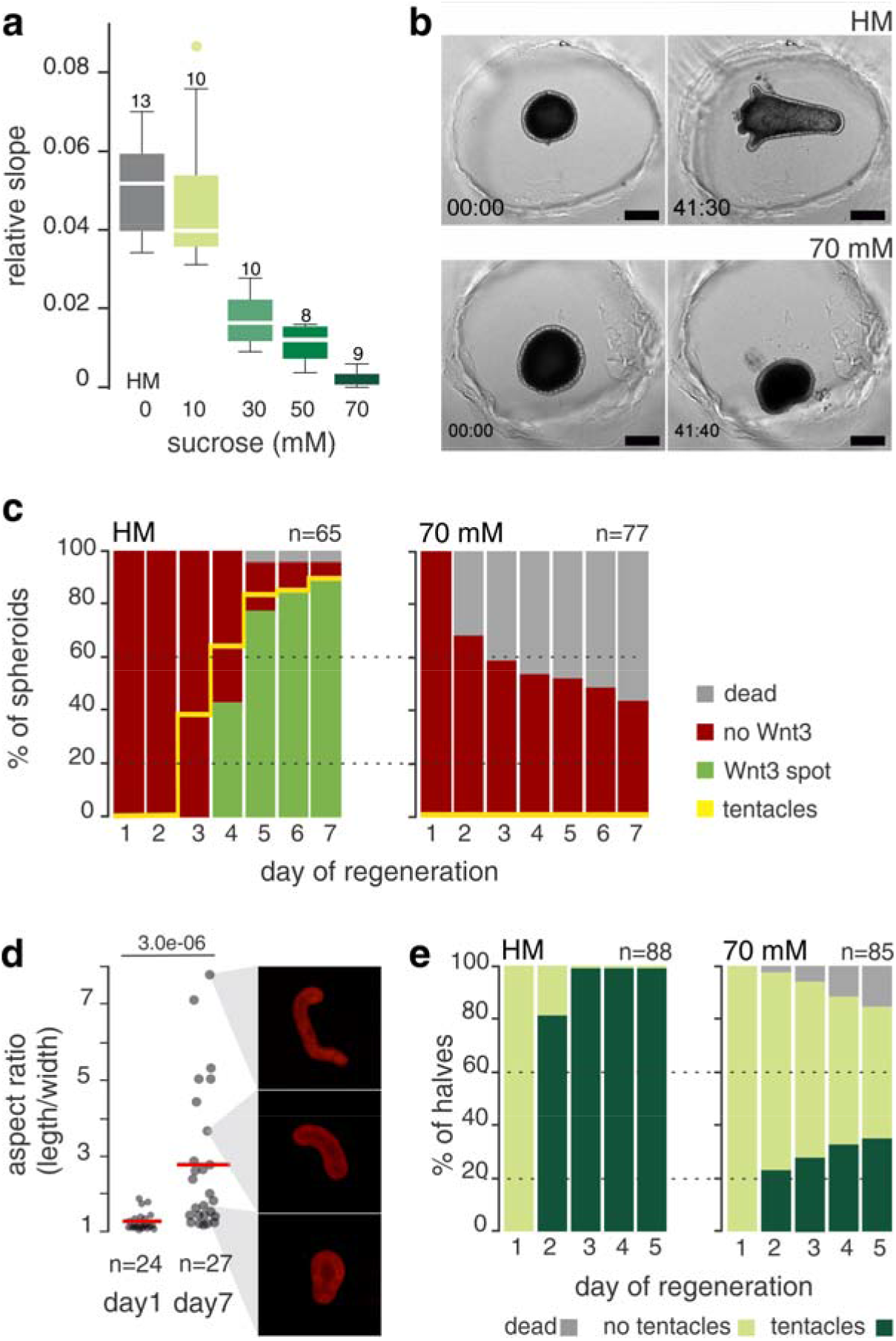
Mechanical oscillations are required for *Hydra* regeneration. (a) Spheroid inflation slope decreases as the osmolarity of the medium increases. The number of samples quantified in each condition is indicated above the boxes. See also Suppl. Fig.2i for details of slope quantification. (b) Snapshots of Suppl. Videos 1 and 2, illustrating the inability of the spheroids to regenerate in isotonic medium. (c) Quantification of morphological regeneration (tentacle appearance) and molecular symmetry breaking (Wnt3 spot) in a *Wnt3::GFP* reporter line spheroids under control (HM) and isotonic conditions (70 mM sucrose). Cumulative plot from n samples collected from 3 independent replicates. (d) The aspect ratio (length/width) of spheroids in isotonic conditions at the beginning and end of the regeneration time course. N=3 independent experiments. P-value from the Wilcoxon rank-sum test is indicated. (e) Head regeneration in the headless halves of bisected animals in control vs. isotonic conditions. Animals were bisected at 50 % body length. A total of n samples from 3 independent replicates were combined.

**Suppl. Fig.2.**
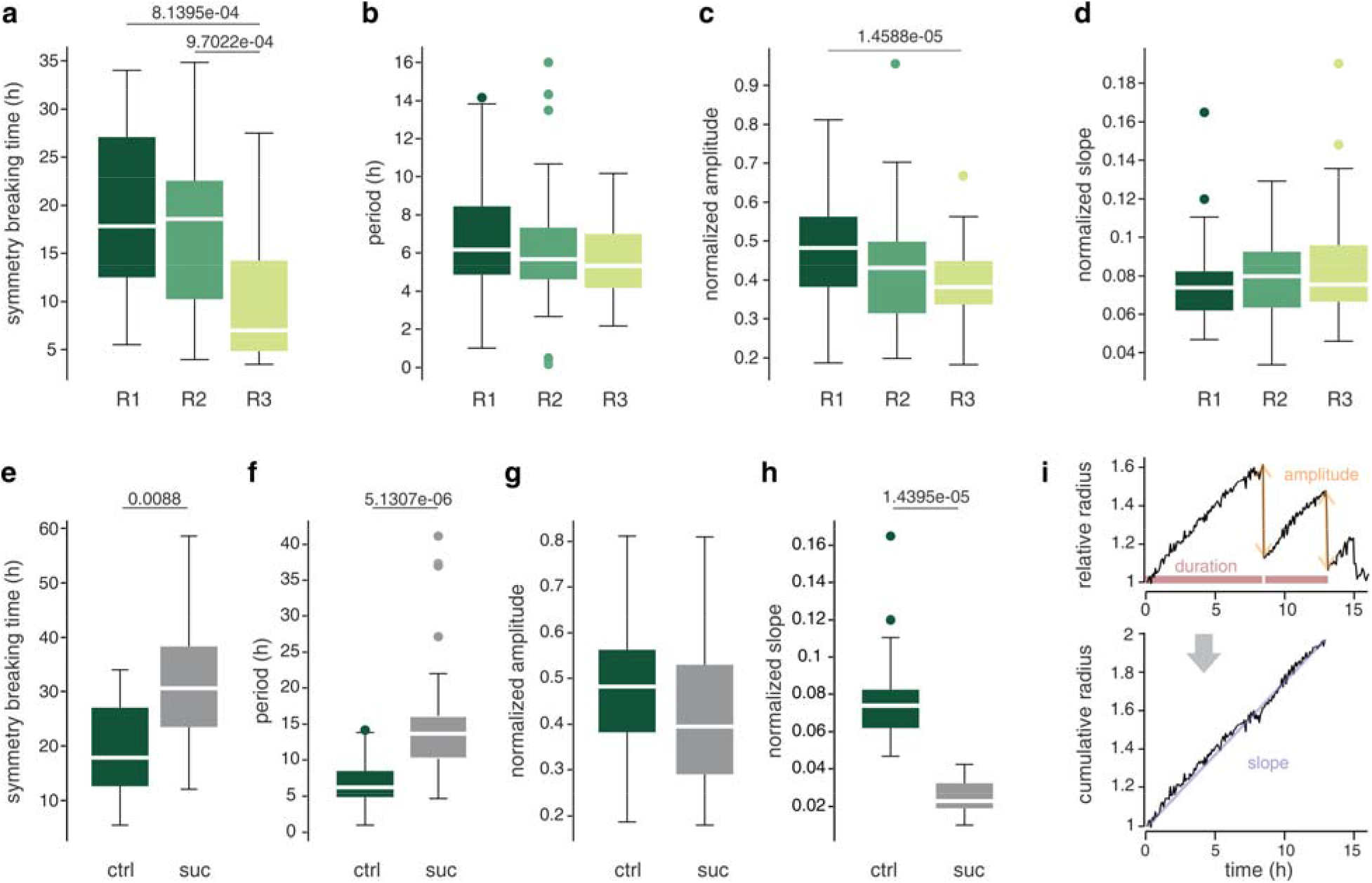
Quantification of mechanical oscillation parameters. (a) Symmetry breaking time, measured as the time of Phase I to Phase II transition for spheroids derived from different axial positions (R1-R3) in control conditions. (b) Period of oscillations for the same samples. (c) Amplitude of oscillations for the same R1, R2 and R3 samples, indicating the maximum size of the spheroid in each cycle before it ruptures. Note the slight axial gradient of amplitudes, which might suggest a gradient in the resistance of the tissue to rupture. However, this gradient alone is not enough to explain the much more pronounced differences in the requirements for mechanical stimulation among the pieces. (d) Slope of inflation. Data shown here (A-D) correspond to the samples in Fig. 2A and are measured on spheroid radius data normalized to the initial size. (e) Comparisons of symmetry breaking time for R1 spheroids in control (HM, for samples in Suppl. Fig.2A) and in 30 mM sucrose. Similar comparisons for oscillation duration (f), oscillation amplitude (g) and inflation slope (h) which, as expected, decreases significantly, resulting in an increase of the oscillation period. (i) An example of the measured parameters. Note that the period and amplitude are quantified for each individual oscillation cycle, while the slope is measured for the entire Phase I, ignoring the deflation events. See (*27*) for details of the quantification procedure. Statistical comparisons were done using the Mood’s median test. Numbers in the panels indicate significant p-values.

**Suppl. Fig.3.**
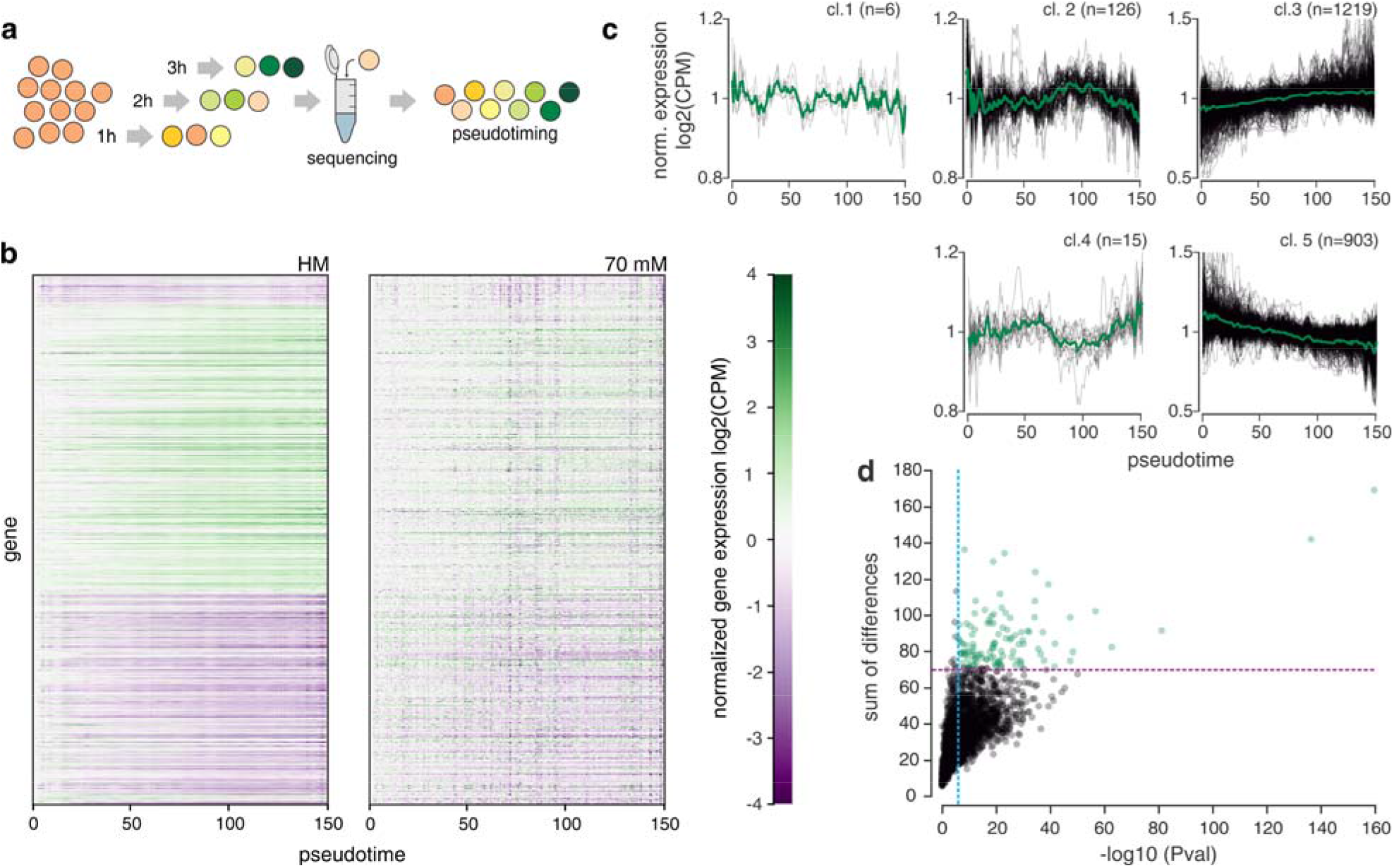
Time series gene expression profiling for spheroids in control (HM) and isotonic (70mM sucrose) conditions. (a) Schematic of the experimental design. Spheroids were left to develop either in HM or isotonic medium after cutting. 8 spheroids were sampled every hour, collected and submitted individually to RNA-sequencing. Transcriptomics data later allowed to reconstruct the temporal development of spheroids (pseudotime) based on the similarity between samples. (b) Heatmaps of the temporal progressions for 2269 genes that change their expression at least 2-fold during the course of normal regeneration. Genes are clustered based on their behavior in control conditions. Note that in the isotonic conditions (70 mM sucrose) some of these genes behave differently (presented in Fig. 3D), while others remain unaffected. In both datasets, the initial log_2_(CPM) values were subtracted for each gene. (c) Plots of the average behavior of the 5 gene clusters identified with hierarchical clustering in the control (HM) sample time-series. Black lines show individual genes, while the thick green line is the cluster average. Number in parentheses indicate the number of genes in each cluster. (d) Selecting the top 10 % mechanosensitive genes. Only genes with a p value < 10^-6^ for temporal changes in the difference between conditions were considered significant (blue line). Purple line indicates the magnitudebased cutoff (sum of differences between the conditions) for the top 10% of significantly changing genes (green dots).

**Suppl. Fig.4.**
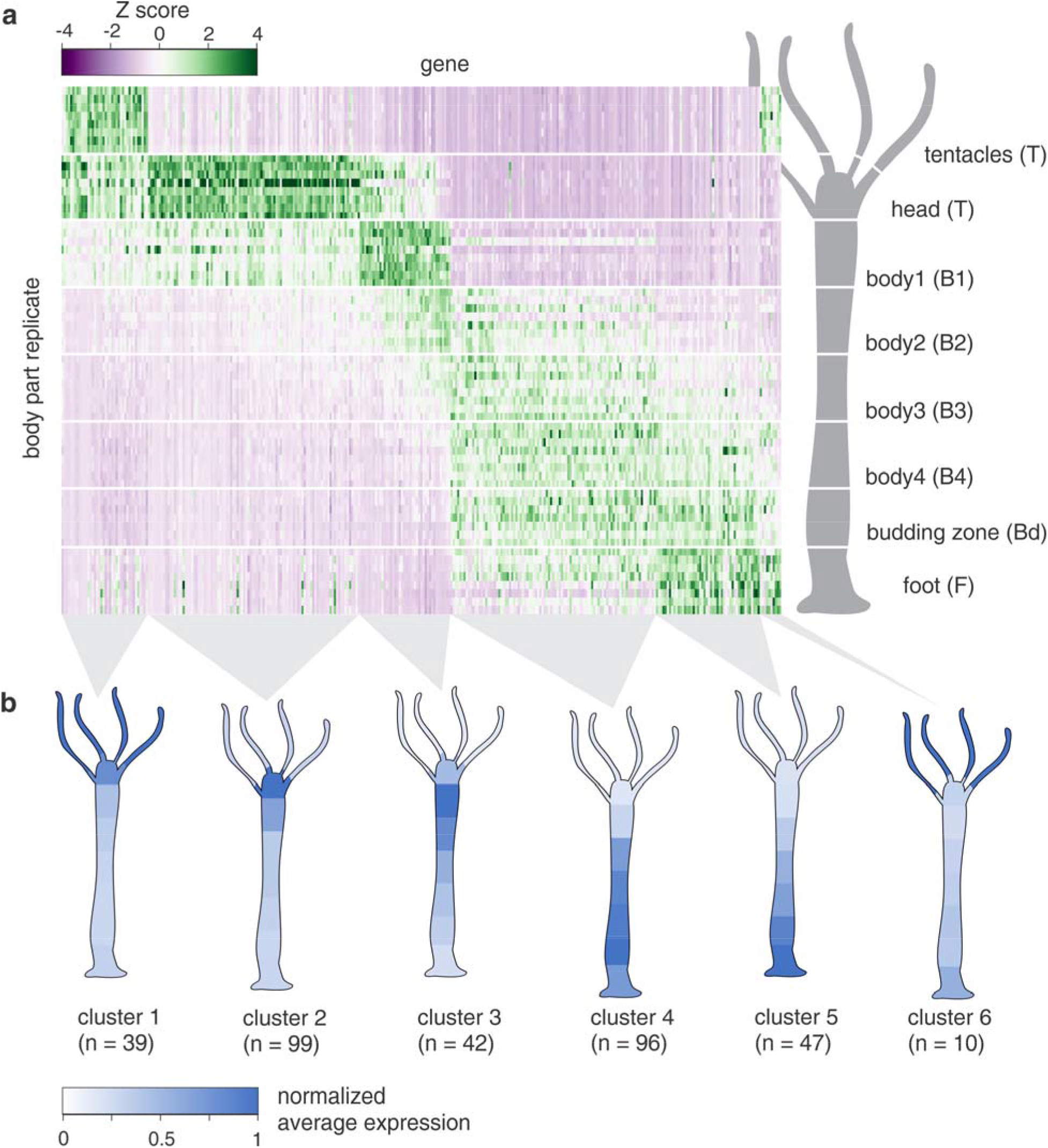
Gene expression profiling along the Hydra Oral/Aboral axis. (a) Heatmap of genes with axially graded expression, identified in the positional RNA-sequencing. Rows correspond to individual replicates for each body position. Color coding indicates the Z-score (standard deviations above or below the mean expression of a gene across all segments after collapsing biological replicates). These data were used to generate the PCA map of different positional identities in Fig 3A-C. (b) Clusters of differentially expressed genes along the O/A axis, identified with k-means clustering. The schematics below show average expression patterns for each cluster, normalized to the body part with highest expression levels.

**Suppl. Fig.5.**
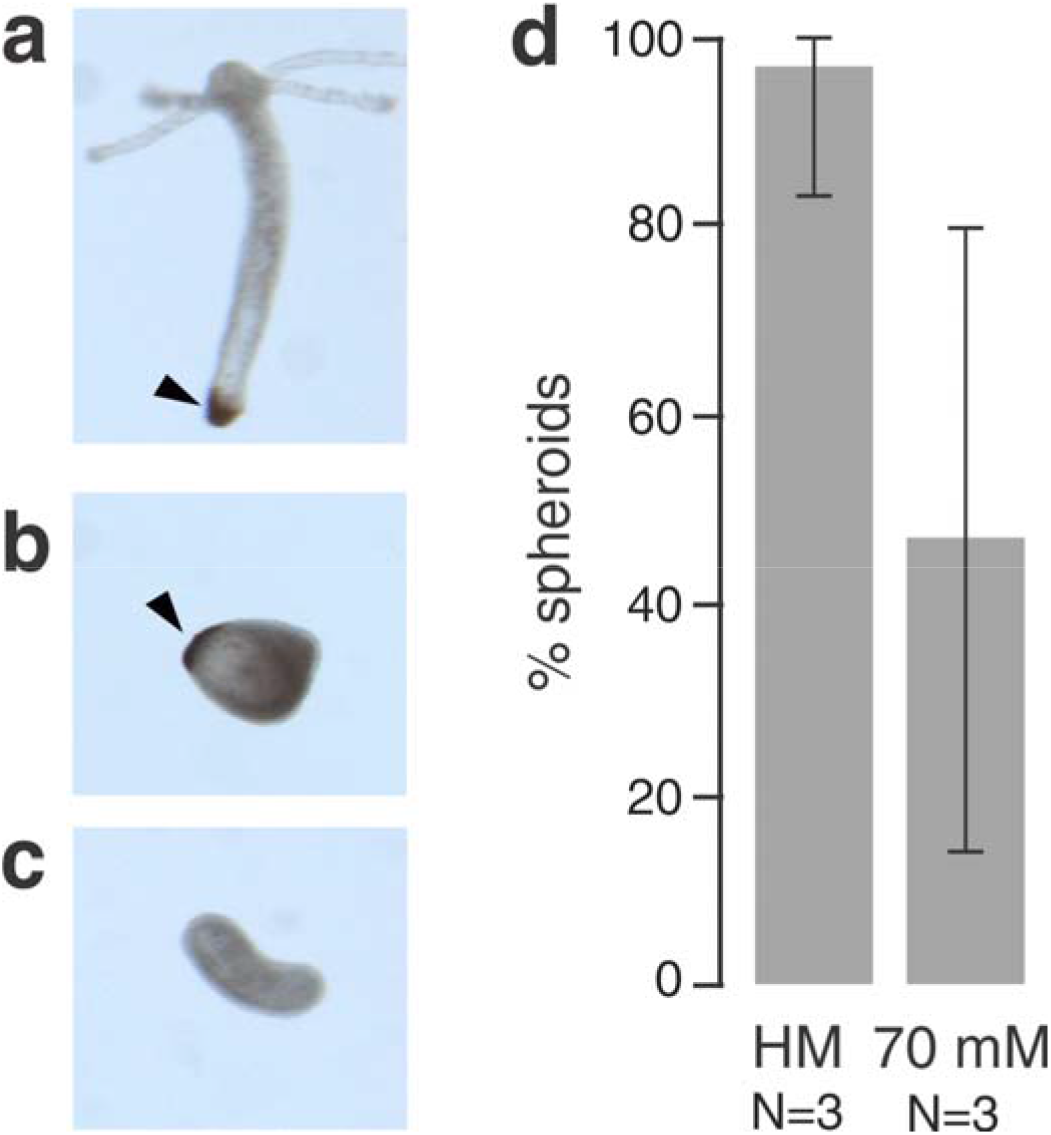
Emergence of the foot organizer in spheroids without mechanical stimulation. (a) Representative spheroid that regenerated under control conditions shows peroxidase staining in the basal disc (foot). (b) Similar foot peroxidase staining was observed in some spheroids cultured in 70 mM sucrose in the absence of a head. (c) Elongated spheroids do not always have a peroxidase marked foot. (d) Quantification of foot marker presence in control (HM) or isotonic medium (70mM sucrose) grown Hydra spheroids. Plot shows the average percentages of peroxidase-positive spheroids from 3 independent experiments. Error bars indicate 95% confidence intervals. 42 spheroids were examined in total for the control, and 26 for isotonic conditions.

**Suppl. Fig.6.**
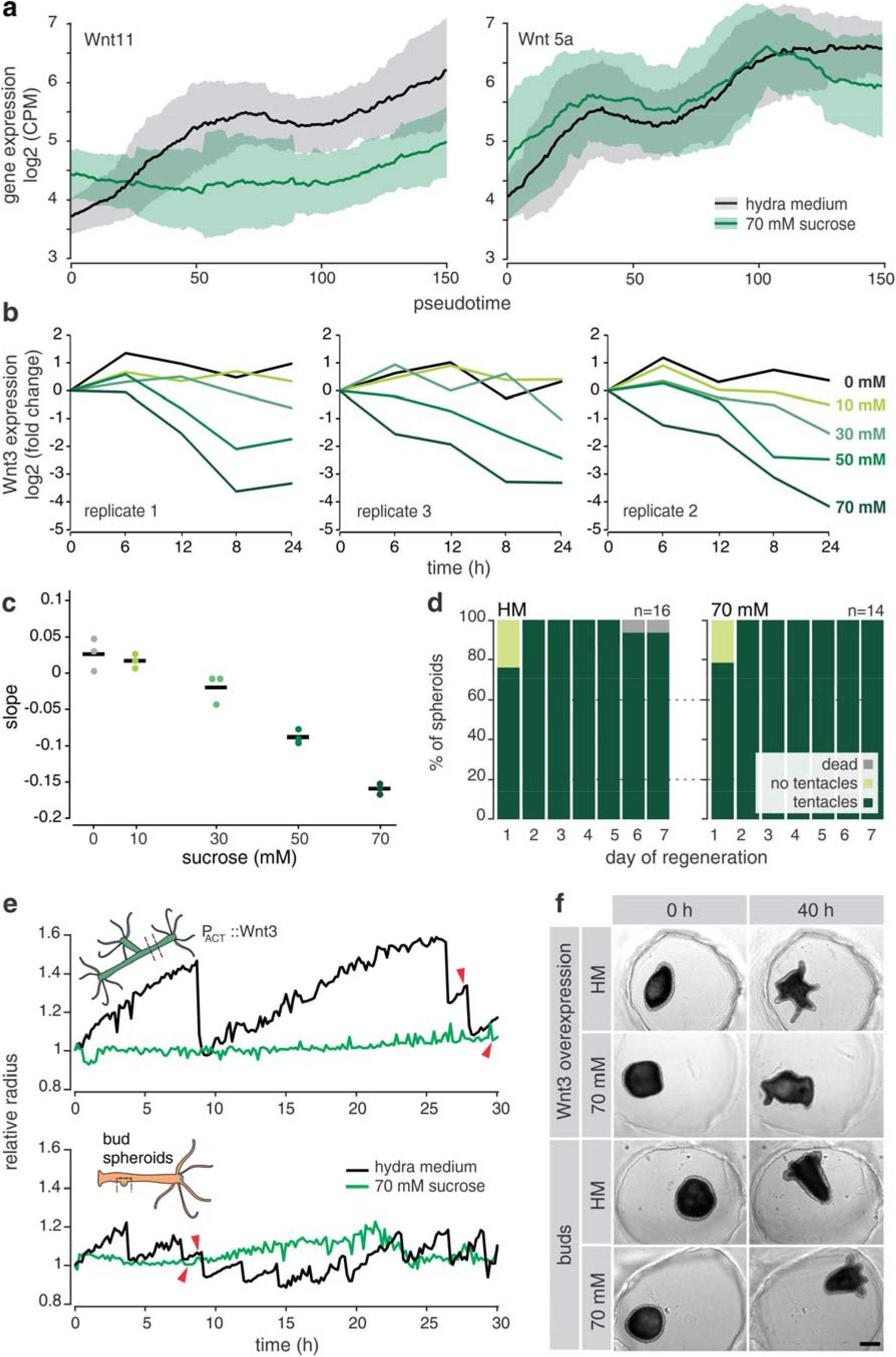
Wnt signaling depends on mechanical stimulation. (a) Differential requirements of mechanical stimulation for the expression of non-canonical Wnt signaling ligands. While the expression pattern of Wnt11 (Wnt/PCP pathway) is sensitive to the removal of mechanical oscillations, Wnt5a (Wnt/Ca^2+^ pathway) does not show changes in the pattern of expression. Lines show a moving mean over the temporal (pseudotime) gene expression data and the shaded area represents moving standard deviation. The size of the moving window is 36 time points. (b) Individual replicates of the qPCR time course from Fig. 4c (c) Slopes of straight lines fitted on the Wnt3 expression data from (b). Dots indicate measurements in individual replicates, and lines represent averages. (d) Spheroids derived from buds regenerate equally well irrespective of the presence or absence of mechanical stimulation. Cumulative plots of n samples combined from 3 independent replicates. (e) Quantifications of the spheroid behavior for Movies S3 – S6 representative of regeneration rescue in the presence of Wnt3. Red arrowheads indicate the emergence of the first tentacle. The top plot shows the radius behavior for Wnt3 overexpressing spheroids. Note the extended Phase I oscillations in HM, which probably reflect changes of tissue mechanical properties in response to high Wnt3 levels. Bud spheroids (lower plot), on the other hand, only show Phase II oscillations. This indicates a possibly established mouth, which is consistent with the early emergence of tentacles. (f) Snapshots of the initial time points and regenerated animals from Suppl. Videos 3 – 6 showing a successful regeneration irrespective of the medium osmolarity. Scale bar, 200 μm. All images are of the same scale.

**Suppl. Fig.7.**
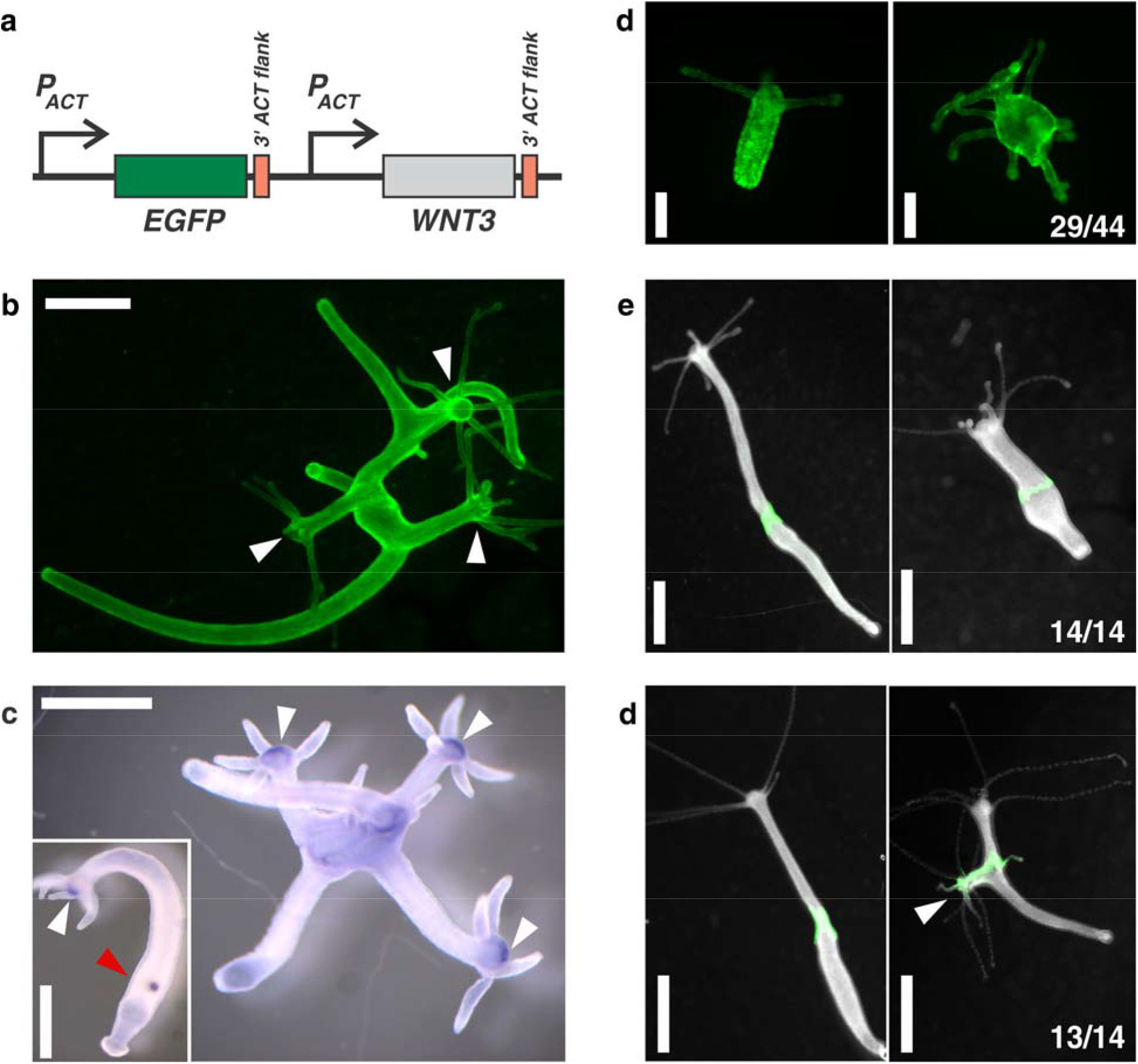
Overexpression of Wnt3 in the ectoderm drives the emergence of new heads. (a) Schematic of the construct used to generate transgenic animals. Both *GFP* and *WNT3* expression is driven by *Hydra* actin promoter. Both genes are also flanked by a 3’ UTR of the *Hydra* actin gene. GFP serves as a transgenesis marker (b) An example of a fully transgenic individual expressing the construct in the ectoderm and showing a typical branched morphology with ectopic heads (arrowheads). (c) Whole mount *in situ* hybridization for *brachyury (Hy_Bra1)* – a head marker and a downstream target of Wnt signaling. Note that event though Wnt3 is expressed uniformly, *brachyury* expression is mostly restricted to the hypostomes (arrowhead). Wild type animal is inset for comparison. Expression can be seen in the hypostome (white arrowhead) as well as in the budding zone (red arrowhead), indicating the onset of budding. (d) Examples of regenerated Wnt3-overexpressing spheroids. While some spheroids regenerate into morphologically normal animals (left), a considerable portion of regenerates shows multiple ectopic tentacles (right) probably reflecting the head-like identity of the tissue conferred by Wnt3. As the animals grow, these tentacles typically cluster into several heads. The numbers indicate the incidence of the multi-tentacle phenotype in regenerating Wnt3-overexpressing spheroids. Data were pooled from three independent experiments. (e) When a piece of tissue from a standard animal is transplanted to a corresponding axial position in another animal, the grafted tissue assimilates within the host body column. Shown is an animal 24h (left) and 5 days (right) after grafting. The graft comes from a GFP-expressing animal. As the numbers indicate, this behavior was observed for all grafts (data pooled from three independent experiments). (f) Unlike in the control situation, when the graft originates from a Wnt3-overexpressing donor, it is able to induce the formation of ectopic heads within the body column (arrowhead). Note that the newly formed head is only partially GFP-positive, indicating the recruitment of host tissue similar to organizer transplantation experiments. This phenotype was observed in all but one animal by day 5 post transplantations (in three independent replicates, pooled counts are given in the figure). In one animal, the graft appeared only to have assimilated by developed and ectopic head later on. Panels E and F show fluorescence images merged with darkfield images of the entire animals. Scale bars in all panels are 1 mm, except for panel D, where scale bars correspond to 200 μm.

## References

Baron, R., and Kneissel, M. (2013). WNT signaling in bone homeostasis and disease: from human mutations to treatments. Nat Med 19, 179–192.

Benos, D.J., Kirk, R.G., Barba, W.P., and Goldner, M.M. (1977). Hyposmotic fluid formation in Hydra. Tissue Cell 9, 11–22.

Bode, H., Lengfeld, T., Hobmayer, B., and Holstein, T.W. (2008). Detection of expression patterns in Hydra pattern formation. Methods Mol Biol 469, 69–84.

Braun, E., and Keren, K. (2018). Hydra Regeneration: Closing the Loop with Mechanical Processes in Morphogenesis. Bioessays 40, e1700204.

Broun, M., Gee, L., Reinhardt, B., and Bode, H.R. (2005). Formation of the head organizer in hydra involves the canonical Wnt pathway. Development 132, 2907–2916.

Cha, B., Geng, X., Mahamud, M.R., Fu, J., Mukherjee, A., Kim, Y., Jho, E.-H., Kim, T.H., Kahn, M.L., Xia, L., et al. (2016). Mechanotransduction activates canonical Wnt/ß-catenin signaling to promote lymphatic vascular patterning and the development of lymphatic and lymphovenous valves. Genes Dev 30, 1454–1469.

Chan, C.J., Costanzo, M., Ruiz-Herrero, T., Mönke, G., Petrie, R.J., Bergert, M., Diz-Muñoz, A., Mahadevan, L., and Hiiragi, T. (2019). Hydraulic control of mammalian embryo size and cell fate. Nature 571, 112–116.

Chera, S., Ghila, L., Dobretz, K., Wenger, Y., Bauer, C., Buzgariu, W., Martinou, J.-C., and Galliot, B. (2009). Apoptotic cells provide an unexpected source of Wnt3 signaling to drive hydra head regeneration. Developmental Cell 17, 279–289.

Ferenc, J. & Tsiairis, C. D. Studying Mechanical Oscillations during Whole Body Regeneration in *Hydra* in *Whole-Body Regeneration*, B. Galliot, S. Blanchoud, Eds. (Methods in Molecular Biology, Springer, 2020), in press

Fernández-Sánchez, M.E., Barbier, S., Whitehead, J., Béalle, G., Michel, A., Latorre-Ossa, H., Rey, C., Fouassier, L., Claperon, A., Brullé, L., et al. (2015). Mechanical induction of the tumorigenic ß-catenin pathway by tumour growth pressure. Nature 523, 92–95.

Hannezo, E., and Heisenberg, C.-P. (2019). Mechanochemical Feedback Loops in Development and Disease. Cell 178, 12–25.

Heisenberg, C.-P., and Bellaïche, Y. (2013). Forces in tissue morphogenesis and patterning. Cell 153, 948–962.

Hobmayer, B., Rentzsch, F., Kuhn, K., Happel, C.M., Laue, von, C.C., Snyder, P., Rothbächer, U., and Holstein, T.W. (2000). WNT signalling molecules act in axis formation in the diploblastic metazoan Hydra. Nature 407, 186–189.

Hoffmeister, S., & Schaller, H. C. (1985). A new biochemical marker for foot-specific cell differentiation in hydra. Wilhelm Roux’s Archives of Developmental Biology 194(8), 453–461.

Kücken, M., Soriano, J., Pullarkat, P.A., Ott, A., and Nicola, E.M. (2008). An osmoregulatory basis for shape oscillations in regenerating hydra. Biophys J 95, 978–985.

Lengfeld, T., Watanabe, H., Simakov, O., Lindgens, D., Gee, L., Law, L., Schmidt, H.A., Özbek, S., Bode, H., and Holstein, T.W. (2009). Multiple Wnts are involved in Hydra organizer formation and regeneration. Dev Biol 330, 186–199.

Li, J., Wang, Z., Chu, Q., Jiang, K., Li, J., and Tang, N. (2018). The Strength of Mechanical Forces Determines the Differentiation of Alveolar Epithelial Cells. Developmental Cell 44, 297–312.e5.

Loh, K.M., van Amerongen, R., and Nusse, R. (2016). Generating Cellular Diversity and Spatial Form: Wnt Signaling and the Evolution of Multicellular Animals. Developmental Cell 38, 643–655.

MacWilliams, H.K. (1983). Hydra transplantation phenomena and the mechanism of Hydra head regeneration. II. Properties of the head activation. Dev Biol 96, 239–257.

Meinhardt, H. (1993). A model for pattern formation of hypostome, tentacles, and foot in hydra: how to form structures close to each other, how to form them at a distance. Dev Biol 157, 321–333.

Nakamura, Y., Tsiairis, C.D., Özbek, S., and Holstein, T.W. (2011). Autoregulatory and repressive inputs localize Hydra Wnt3 to the head organizer. Proc. Natl. Acad. Sci. U.S.A. 108, 9137–9142.

Newman, S.A., Forgacs, G., and Muller, G.B. (2006). Before programs: the physical origination of multicellular forms. Int J Dev Biol 50, 289–299.

Picelli, S., Björklund, Å.K., Reinius, B., Sagasser, S., Winberg, G., and Sandberg, R. (2014a). Tn5 transposase and tagmentation procedures for massively scaled sequencing projects. Genome Res 24, 2033–2040.

Picelli, S., Faridani, O.R., Björklund, Å.K., Winberg, G., Sagasser, S., and Sandberg, R. (2014b). Full-length RNA-seq from single cells using Smart-seq2. Nat Protoc 9, 171–181.

Pukhlyakova, E., Aman, A.J., Elsayad, K., and Technau, U. (2018). ß-Catenin-dependent mechanotransduction dates back to the common ancestor of Cnidaria and Bilateria. Proc. Natl. Acad. Sci. U.S.a. 115, 6231–6236.

Ruiz-Herrero, T., Alessandri, K., Gurchenkov, B.V., Nassoy, P., and Mahadevan, L. (2017). Organ size control via hydraulically gated oscillations. Development 144, 4422–4427.

Ryan, A.Q., Chan, C.J., Graner, F., and Hiiragi, T. (2019). Lumen Expansion Facilitates Epiblast-Primitive Endoderm Fate Specification during Mouse Blastocyst Formation. Developmental Cell 51, 684–697.e684.

Schindelin, J., Arganda-Carreras, I., Frise, E., Kaynig, V., Longair, M., Pietzsch, T., Preibisch, S., Rueden, C., Saalfeld, S., Schmid, B., et al. (2012). Fiji: an open-source platform for biological-image analysis. Nat Methods 9, 676–682.

Shahbazi, M.N. (2020). Mechanisms of human embryo development: from cell fate to tissue shape and back. Development 147, dev190629.

Shimizu, H. (2012). Transplantation analysis of developmental mechanisms in Hydra. Int J Dev Biol 56, 463–472.

Shinozuka, T., Takada, R., Yoshida, S., Yonemura, S., and Takada, S. (2019). Wnt produced by stretched roof-plate cells is required for the promotion of cell proliferation around the central canal of the spinal cord. Development 146, dev159343.

Soriano, J., Rüdiger, S., Pullarkat, P., and Ott, A. (2009). Mechanogenetic coupling of Hydra symmetry breaking and driven Turing instability model. Biophys J 96, 1649–1660.

Teague, B.P., Guye, P., and Weiss, R. (2016). Synthetic Morphogenesis. Cold Spring Harb Perspect Biol 8, a023929.

Vogg, M.C., Beccari, L., Iglesias Ollé, L., Rampon, C., Vriz, S., Perruchoud, C., Wenger, Y., and Galliot, B. (2019a). An evolutionarily-conserved Wnt3/ß-catenin/Sp5 feedback loop restricts head organizer activity in Hydra. Nature Communications 10, 312–315.

Vogg, M.C., Galliot, B., and Tsiairis, C.D. (2019b). Model systems for regeneration: Hydra. Development 146, dev177212.

Wittlieb, J., Khalturin, K., Lohmann, J.U., Anton-Erxleben, F., and Bosch, T.C.G. (2006). Transgenic Hydra allow in vivo tracking of individual stem cells during morphogenesis. Proc Natl Acad Sci USA 103, 6208–6211.

